# Geochemical constraints on bacteriophage infectivity in terrestrial environments

**DOI:** 10.1101/2023.04.10.536276

**Authors:** Hans K. Carlson, Denish Piya, Madeline L. Moore, Roniya Thapa Magar, Nathalie H. Elisabeth, Adam M. Deutschbauer, Adam P. Arkin, Vivek K. Mutalik

**Author notes:** to whom correspondence should be addressed Hans K. Carlson, Vivek K. Mutalik.

## Abstract

Lytic phages can be potent and selective inhibitors of microbial growth and can have profound impacts on microbiome composition and function. However, there is uncertainty about the biogeochemical conditions under which phage predation can proceed and modulate microbial ecosystem function, particularly in terrestrial systems. Ionic strength is known to be critical for infection of bacteria by many phages, but there is limited quantitative data on ion thresholds for phage infection that can be compared with environmental ion concentrations. Similarly, while carbon composition varies in terrestrial environments, we know little of which carbon sources favor or disfavor phage infection and how these higher order interactions impact microbiome function. Here, we measured the half-maximal effective concentrations (EC_50_) of 80 different inorganic ions for the infection of *E. coli* with two canonical dsDNA and ssRNA phages, T4 and MS2, respectively. We found that many alkaline earth metals and alkali metals enabled successful lytic infection but that the ionic strength thresholds varied for different ions between phages. Additionally, using a freshwater nitrate reducing microbiome, we found that the ability of lytic phage to influence nitrate reduction end-products was dependent on the carbon source as well as the ion concentration. For all phage:host pairs we tested, the ion EC_50_s for phage infection we measured exceed the ion concentrations found in many terrestrial freshwater systems. Thus, our findings support a model where the influence of phages on terrestrial microbial functional ecology is greatest in hot spots and hot moments such as metazoan guts, drought influenced soils, or biofilms where ion concentration is locally or transiently elevated and carbon source composition is of a sufficiently low complexity to enrich for a dominant phage susceptible population.

**Significance:** Viral-prokaryote dynamics greatly influence microbial ecology and the earth’s biogeochemical cycles. Thus, identifying the key environmental controls on phage predation is critical for predictive microbial ecology. Here we conduct laboratory experiments that implicate ionic strength and carbon composition as major controls on phage interactions with bacterial hosts in terrestrial microbiomes. We propose a model in which terrestrial phage predation is most favored in drought impacted soils and in higher ionic strength environments such as metazoan guts or between adjacent cells in biofilms.

## Introduction

Bacteriophages are ubiquitous in natural microbial communities and can play important roles in modulating microbial element cycling [1–7]. However, we lack robust mechanistic models to enable predictions of phage-host dynamics and ecological interactions, particularly in complex environments such as soils and sediments. While some studies have attempted to demonstrate an impact of phages on soil element cycling, results are variable and apparently context-dependent [1–5]. As such, the magnitude of the impact of phage predation on element cycling across terrestrial ecosystems and the mechanistic biogeochemistry of this process remain elusive.

In marine environments, phage lysis is responsible for ∼40-50% of all bacterial mortality, approximately equivalent to the contribution of protozoal predation [6]. Carbon released from dead bacterial cells following phage predation drives a “viral shunt” in the marine carbon cycle [7]. In terrestrial systems, however, the contribution of phage to element cycling dynamics is much less well characterized and appears to be more variable, particularly in extremely heterogeneous environments such as soils [1–5].

In the absence of a mechanistic model of phage dynamics in terrestrial ecosystems it is difficult to interpret correlations between bacterial:viral ratios and environmental parameters to identify causal relationships [8]. Several studies have identified strong relationships between host bacterial blooms and predatory phage in natural systems [9, 10]. It thus follows that environmental parameters which influence host abundance are important for predatory phage success. For example, phage abundance can be influenced by the presence of essential microbial nutrients and energy sources required for host growth [11, 12] with the virus to microbial ratio increasing with microbial cell density [12]. Thus, in soils, because particles of detrital carbon or root exudates have higher microbial densities these should be hotspots of phage predation. Furthermore, the quality of carbon will determine the composition of the heterotrophic microbial population [13, 14]. It follows that as carbon composition shifts, the abundance of phage susceptible microbial populations will also shift to alter the possible impact of phages on microbial element cycling processes.

Inorganic ion gradients are also likely to play a major role in phage:host interactions in terrestrial ecosystems. It has been known for nearly a century that certain alkali metals and alkaline earth metals including calcium, sodium and magnesium are important for phage infectivity [15, 16]. This is in part because these cations interact with phage tail fibers to disaggregate them and enable phage binding to host cell surface receptors [17, 18]. Cations are also essential to neutralize negatively charged membranes and phage to enable phage binding [19, 20]. Most phages and bacteria are negatively charged under circumneutral environmental conditions [21]. For model phage:host pairs, greater than 1 mM Ca^2+^ or Mg^2+^ or greater than 10-40 mM Na^+^ was required for efficient phage binding [19]. Cations are also important for phage retention on soil and sediment particles via similar charge neutralization mechanisms [20, 22, 23]. Membrane fluidity is also influenced by ionic strength [24] and this may alter phage receptor binding, particularly for phages that bind to outer membrane lipids. Also, the persistence length of DNA packaged in phages is very sensitive to changes in ionic strength with more disordered DNA at lower ionic strength likely hampering infectivity [25–27]. Finally, the expression of some phage receptors are induced at higher ionic strength. The T4 phage receptor, OmpC is upregulated at ionic strengths between 10-500 mM [28] and the MS2 receptor, the type 1 conjugative pilus, is similarly upregulated with increasing ionic strength [29]. Thus, there are many indications that there are environmentally relevant ion thresholds below which phage predation will have a decreased impact on microbiomes. We anticipated that identifying these thresholds in well controlled laboratory systems would provide valuable constraints for models of environmental microbial ecology.

Measuring higher order interactions between geochemical parameters and microbial ecology requires high-throughput approaches to simulate and recapitulate environmental conditions under controlled laboratory settings [14, 30, 31]. Bacteria and phages can persist across broad ranges of ion and carbon complexity and concentration, and some studies have observed correlations between viral:host dynamics and environmental parameters [8]. However we lack detailed mechanistic understanding of how complex natural environmental gradients of carbon and inorganic ions influence phage predation. Furthermore, the use of non-standard media in previous laboratory low-throughput and low-resolution studies [15–19, 32] make systematic comparisons challenging.

In this study, we hypothesized that the nature and concentration of ions and carbon sources in a microbial ecosystem are major controls on phage-host dynamics and designed laboratory experiments to measure these influences on model phage:host interactions. We used the model interaction between T4 and MS2 phages and *E. coli* hosts to quantify inorganic ion thresholds on phage infectivity. We also used a cultured nitrate reducing microbiome from freshwater aquatic sediment to study the higher order interactions between carbon composition, inorganic ion concentration and lytic phage in controlling microbial element cycling. We found that microbial community composition and respiratory activity is sensitive to changes in the composition and concentration of both carbon sources and inorganic ions. Specifically, we observed that phages can limit the growth and metabolic activity of populations carrying out a dominant element cycling function (dissimilatory nitrate reduction to ammonium, DNRA), but that phage predation is dependent upon both having a carbon source that can be utilized by the DNRA population and inorganic ion concentrations above that observed in most terrestrial freshwater environments. Therefore, we propose a mechanistic model in which phage predation in terrestrial systems occurs predominantly in hotspots where there is sufficiently low complexity carbon and inorganic ions are locally or transiently concentrated.

## Results

### Influence of inorganic ions and phage on the growth of model E. coli strains

To quantify the influence of inorganic ions on phage:host interactions, we first measured the growth of two model *E. coli* strains, BW25113 (the host for the dsDNA phage T4) and C3000 (the host for the ssRNA phage MS2) in the presence of an array of 80 different serially-diluted inorganic ions using a high-throughput assay platform [30, 31]. We then measured the growth of these host strains across the inorganic ion array in the presence of their corresponding lytic phages at a multiplicity of infection (MOI) of 1 phage per bacterial cell (**Figure 1A**). Importantly, we used an ion-depleted chemically defined medium for our control cultures containing ∼30 mM Na^+^ and sub-millimolar concentrations of other trace minerals and ions. In this ion-depleted media, *E. coli* growth was not inhibited by the addition of lytic phage. Thus, we were able to observe that many inorganic ions displayed similar inhibitory potencies (IC_50_) in the presence and absence of phages (**Table S1**). However, several alkali earth metals and alkali metals displayed a lower half-maximal inhibitory concentration in the presence of phage (**Figure 1B,1C**). These lower inhibitory potencies measured in the presence of phage represent the effective concentration required for phage infection (EC_50_). We also noticed a subtle, but reproducible increase in the IC_50_ of Al^3+^ in the presence of T4 phage (**Figure 1B, 1C**). These results may suggest that phages can protect bacteria from metal toxicity in environments with high concentrations of transition metals.

**Figure 1.**
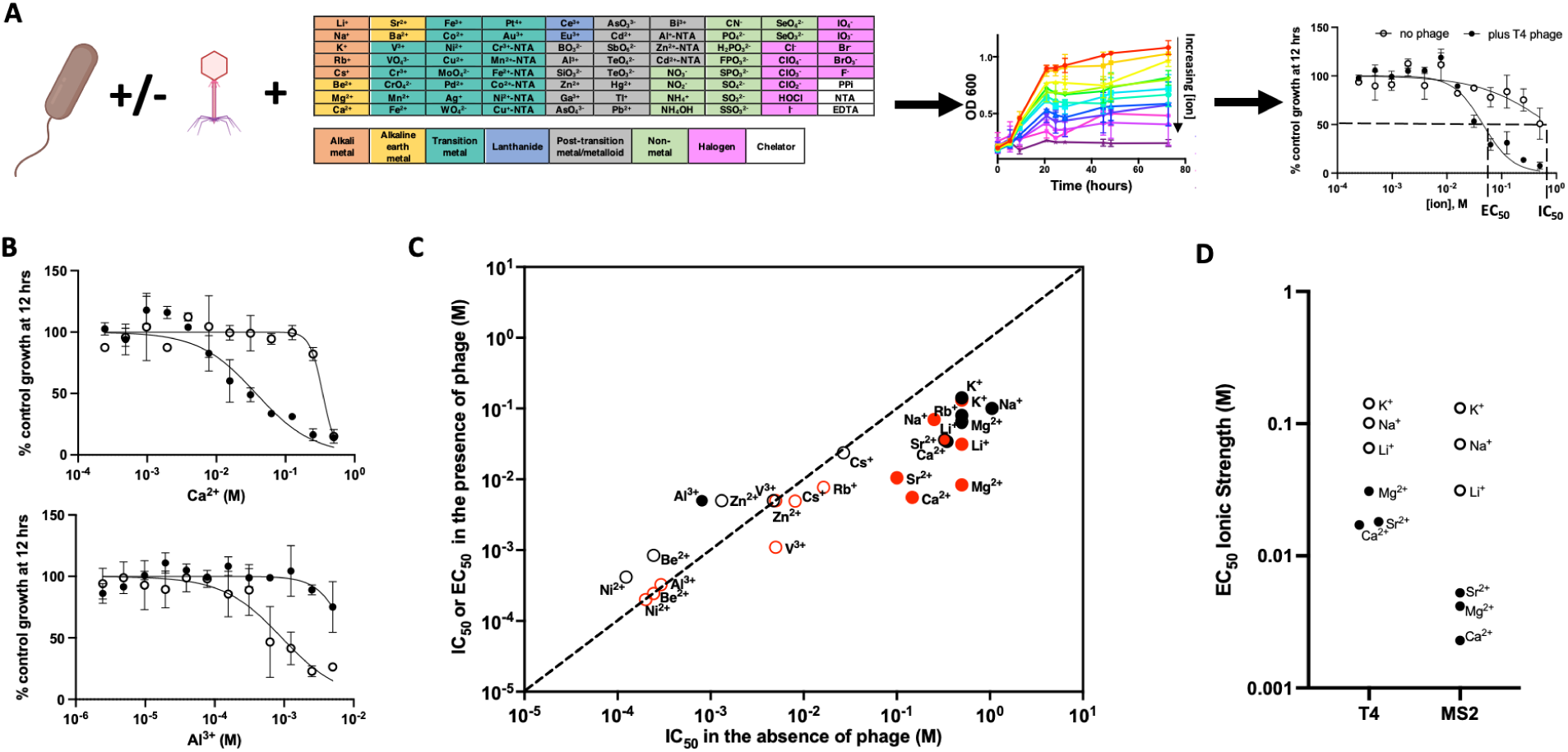
Measurement of concentrations of inorganic ions necessary for lytic phage infection (EC_50_) or direct ion toxicity against *E. coli* (IC_50_). **A.** To measure inorganic ion influence on phage predation bacterial hosts are grown in an ion-depleted chemically defined medium in the presence and absence of phage and in the presence and absence of an array of inorganic ions serially diluted in microplates. Half maximal inhibitory concentrations are quantified in the presence and absence of phage to identify ion toxicity thresholds (IC_50_) and ion thresholds required for phage infection (EC_50_). **B.** Growth of *E. coli* BW25113 cultures at 12 hours as measured by optical density (OD 600) in the presence (closed symbols) or absence (open symbols) of T4 phage and varying concentrations of Ca^2+^ and Al^3+^ relative to control cultures lacking phage or additional ions. **C.** Comparison of selected ion IC_50_ or phage-ion EC_50_ for *E. coli* BW25113 and phage T4 (black), and for *E. coli* C3000 and phage MS2 (red). Closed symbols indicate a significant difference between IC_50_/EC_50_ in the absence/presence of phage. **D.** EC_50_ expressed as ionic strength for ions that enable lytic phage infection (open symbols = alkali metals, closed symbols = alkali earth metals).

Our dose-response assays indicate that the major divalent alkali earth metals (Ca^2+^ and Mg^2+^) and monovalent alkali metals (K^+^ and Na^+^) found in natural waters enable lytic infection by T4 and MS2 (**Figure 1C, 1D**). Of the less common ions in natural waters, Sr^2+^ and Li^+^ enabled efficient phage infection, while Be^2+^, Rb^+^, and Cs^+^ were more toxic with similar inhibitory potencies in the presence and absence of phage. Cs^+^ and Rb^+^ likely interfere with K^+^ metabolism [33][34] while Be^2+^ has the smallest ionic radius and most negative electron affinity of the alkali metal/alkali earth metals we tested. In general, MS2 phage required a lower ionic strength than was required for T4 infection (**Figure 1C, 1D**). Of the ions that enable lytic phage infection, the monovalent ions required higher concentrations even when correcting for the difference in ionic strength conferred from the divalent ions (**Figure 1D**). For example, Ca^2+^ EC_50_ were significantly lower than Na^+^ EC_50_ for both MS2 and T4. Others have reported both specific ion requirements and non-specific ion requirements for phages [35–41]. Compared to these previous studies, however, our results and methods show clear ion specificity across a wider range of inorganic species and concentration ranges. As such, our results demonstrate that phages differ in their ionic strength requirements and that for both MS2 and T4 phages the nature of the inorganic ion contributing to ionic strength influences the concentration of ion necessary for lytic infection. Only a small subset of ions (Ca^2+^, Mg^2+^, K^+^, Na^+^, Sr^2+^, Li^+^) enable lytic phage infection at concentrations lower than the direct ion toxicity to cells.

### Influence of carbon composition and ionic strength on lytic phage modulation of microbial element cycling

We hypothesized that ionic strength also controls phage:host interactions and microbial element cycling in more complex microbiomes. To test this hypothesis we used a model nitrate reducing microbial enrichment culture from aquatic freshwater sediment (FN enrichment culture). This cultured microbiome has diverse heterotrophic bacteria with distinct and varying respiratory capabilities including both denitrifiers and bacteria capable of dissimilatory nitrate reduction to ammonium (DNRA). Previously, we have used this microbiome to demonstrate that specific carbon sources can influence the end-products of microbial nitrate reduction by selectively enriching for microbial strains with particular respiratory enzymatic capabilities [14]. From our previous work [14], we have pure culture isolates from this enrichment including an *Escherichia* and two *Citrobacter* strains that depending on the primary electron donor/carbon source can dominate in catalyzing DNRA. To assess the phage influence on DNRA microbiome structure and function, we isolated and characterized dsDNA phages that can prey on either the *Escherichia* or the two *Citrobacter* isolates (**Figure S1**).

Next, we formulated a cocktail of three phage isolates each capable of lytic infection of one of the dominant DNRA organisms in the enrichment, two *Citrobacter* and one *Escherichia,* and measured the impact of this phage cocktail on ammonium production and strain composition of the enrichment culture (**Figure 2**, **Figure S2**). We tested the addition of the phage cocktail on the enrichment culture recovered on different carbon sources/electron donors (**Figure 2A**) and found that the impact of phage was greater for some carbon sources than others (**Figure 2B, 2C**). For carbon sources in which the *Escherichia* or *Citrobacter* strains dominate, such as D-trehalose, D-glucose or D,L-lactate, the ammonium production was significantly decreased in the presence of the lytic phage cocktail (ANOVA, p<0.05) (**Figure 2B**, **Figure S2**). This corresponded to a subtle but significant decrease in the relative abundance of at least one of the dominant DNRA organisms. In contrast, cultures grown in D-cellobiose and L-sorbose enriched for a *Klebsiella* which is only capable of nitrate reduction to nitrite. Neither of the DNRA capable *Citrobacter* and *Escherichia* utilize D-cellobiose or L-sorbose and as such ammonium accumulation was already low in the absence of phages and the addition of the phage cocktails had little effect on ammonium production. Finally, ethanol and formate enriched for a *Sulfurospirillum* that is capable of DNRA. In these cultures, ammonium accumulation was largely driven by a strain not targeted by the lytic phage cocktail and hence was unaffected by the phage addition. Together, these results demonstrate that the impact of phages on microbial element cycling depends greatly on the composition of the microbiome, which is in turn controlled by changes in nutrients such as electron donors and carbon sources.

**Figure 2.**
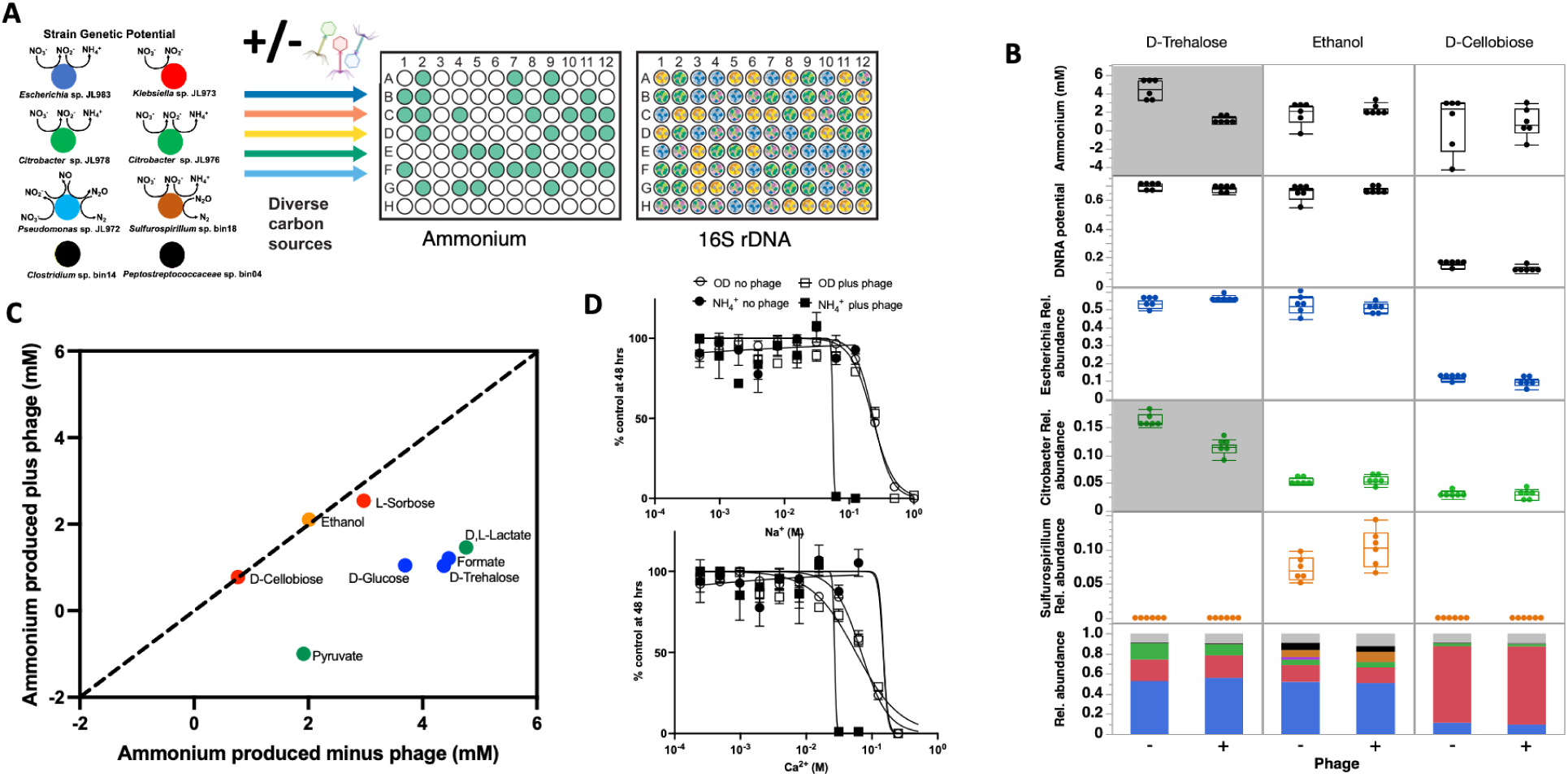
Influence of carbon sources and inorganic ions on a phage cocktail formulated to inhibit the DNRA (dissimilatory nitrate reduction to ammonium) function in a model freshwater nitrate respiring microbiome (FN). **A**. A genomically-characterized nitrate-reducing microbiome is recovered in the presence of diverse carbon sources in the presence and absence of a phage cocktail formulated to target DNRA function in the microbiome. Ammonium and community composition are measured to determine the influence of each carbon source on phage modulation of the microbial element cycling function. **B.** Ammonium production, DNRA genetic potential and relative abundances of dominant strains in the FN microbiome on selected carbon sources. Box and whiskers represent interquartile range. Shaded panels indicate significant differences (ANOVA, p<0.05) between plus (+) and minus (-) phage conditions. In stacked bar plot: Dark Blue = Escherichia, Green = Citrobacter, Red = Klebsiella, Orange = Sulfurospirillum, Light Blue = Pseudomonas, Black = Gram-positive fermenters, Gray = other strains. **C**. Ammonium production by the FN microbiome grown on different carbon sources in the presence and absence of phage cocktail. Colors of points reflect the dominant strain enriched on each carbon source and match color scheme in Panel A. **D**. Dose-response assays to assess inorganic ion requirements for the phage cocktail to inhibit growth (OD 600) or ammonium production in the FN microbiome relative to ion depleted control cultures.

To test whether ionic strength influences the ability of our lytic phage cocktail to modulate the DNRA function of the freshwater microbial enrichment culture, we grew the enrichment on D-trehalose in a low ionic strength medium (∼40 mM Na^+^) with varying concentrations of Na^+^ or Ca^2+^ (**Figure 2D**). D-trehalose is produced by diverse soil microorganisms, including *Enterobacteria*, as an osmoprotectant under drought conditions [42]. Thus, this experiment may simulate the physiological states experienced by *Enterobacteria* in soils as water content and ionic strength varies. Ammonium concentrations in D-trehalose cultures were also extremely responsive to addition of the lytic phage cocktail. We measured both growth and ammonium accumulation by these cultures as a function of Na^+^ or Ca^2+^ concentration. We found that both ions inhibit ammonium accumulation in the presence of phage at lower concentrations than in the absence of phage, demonstrating their importance for lytic phage infection in this model freshwater microbiome. Furthermore, the EC_50_ of these ions against the FN culture were similar as those measured for MS2 or T4 (**Figure 1**).

### Comparison of ion concentrations necessary for phage infection with environmental ion concentrations

Having measured ion EC_50_ thresholds for lytic phage infection of bacterial hosts, we wondered if these concentrations are attained in terrestrial freshwaters. Na^+^, Ca^2+^ and Mg^2+^ concentrations in freshwaters vary by over three orders of magnitude from low ionic strength spring waters [43] to high ionic strength agricultural drainage waters [44] (**Figure 3**), but ion concentrations are much higher in bacterial cytoplasm [45], the human gut or seawater [46]. For Na^+^, none of the freshwaters reached the necessary ion concentrations required for lytic phage infection we measured (**Figure 3A**), but Na^+^ concentrations in the bacterial cytoplasm, human gut or seawater exceed the thresholds for phage infection that we measured. For both Ca^2+^ and Mg^2+^, a few of the extremely high ionic strength agricultural waters from tile drained soils reached the ion concentrations required for MS2 infection, but not the concentrations required for T4 or the freshwater nitrate reducing enrichment culture phage cocktail (**Figure 3B, 3C**). In seawater, Ca^2+^ and Mg^2+^ concentrations exceeded the MS2 phage EC_50_ and in the bacterial cytoplasm Mg^2+^ concentrations exceeded the MS2 phage EC_50_.

**Figure 3.**
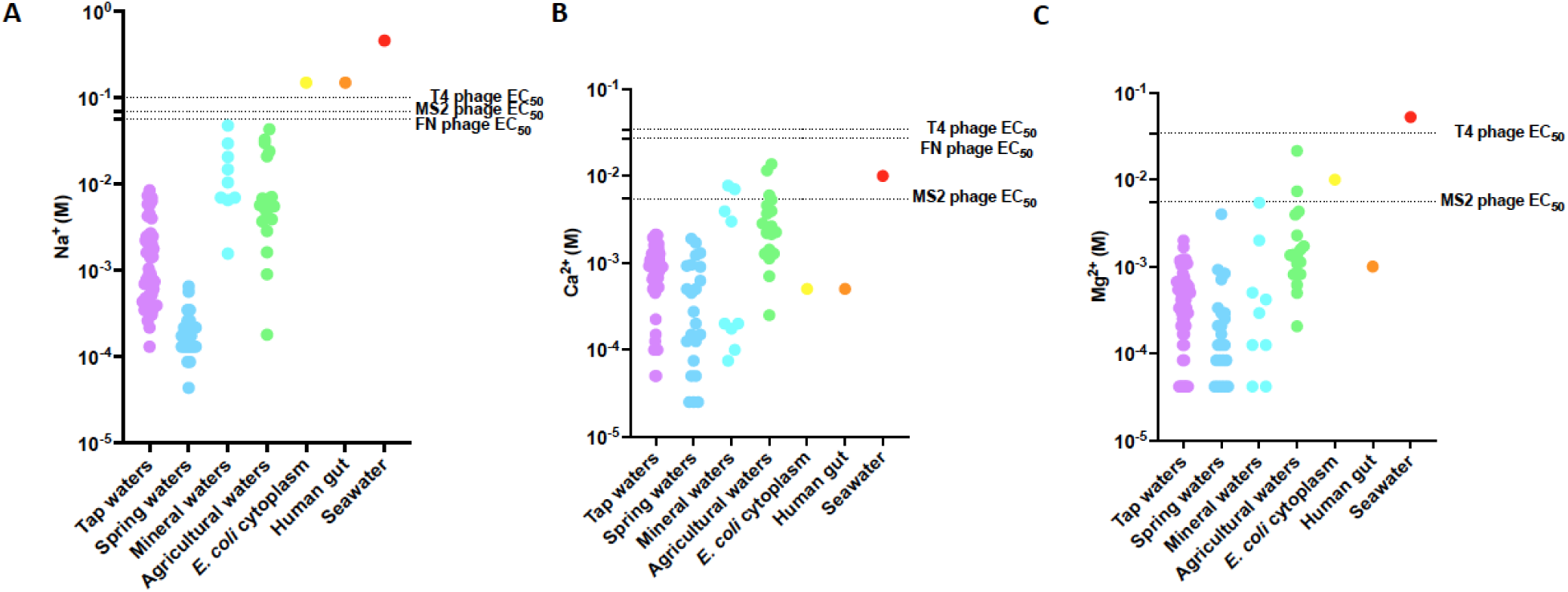
Comparison of ion EC_50_ values for MS2, T4 and freshwater nitrate respiring microbiome (FN) phage cocktail with concentrations of the major ions in diverse freshwaters, the bacterial cytoplasm the human gut or seawater **A**. Na^+^, **B**. Ca^2+^, and **C**. Mg^2+^. (Please see the main text for references).

## Discussion

Overall, our results along with those in the literature lead us to propose that the low ionic strength in many freshwater environments may limit successful lytic phage infection for the Enterobacterial phage host:pairs we studied. In contrast, in metazoan guts, in biofilms adjacent to recently lysed cells or in estuarine/marine environments, ionic strength is not a limitation on these phage:host interactions. Our proposal is supported by ion EC_50_ measured for other soil and freshwater environmental phages of *Bacillus* and *Rhodopseudomonas* genera [39–41]. For example, *Bacillus* phage were shown to require cations above that found in many water systems (∼1-10 mM Ca^2+^ or Mg^2+^, ∼50-100 mM Na^+^) for maximal infectivity [41, 47]. And phages of the model freshwater bacterium, *Caulobacter*, have been shown to require Mg^2+^ at ∼ 1 mM [48, 49]. A recent review of diverse phage studies concluded that on average 2.38 mM Ca^2+^, 5.08 mM Mg^2+^ or ∼100 mM Na^+^ were required for efficient infection [32]. All of these measurements are consistent with the classical theoretical model that neutralization of membrane charge is required for phage binding and that this takes place >1mM for divalent cations (e.g. Ca^2+^ or Mg^2+^) and >10mM for monovalent cations (e.g. Na^+^) [19, 20]. Importantly, these concentrations are higher than are found in many freshwaters (**Figure 3**). In fact the global mean Ca^2+^ concentration in freshwaters is ∼0.1 mM and decreasing due to anthropogenic acidification [50]. Notably, while most phage lysis media rely on high concentrations of the likely non-physiological ions Ca^2+^ or Mg^2+^, our data suggests that Na^+^ may actually be a more important ion for enabling phage infection in many freshwaters because Ca^2+^ or Mg^2+^ concentrations are often too low to support lytic phage infection. Future work should focus on measuring ion thresholds for more phage using more standardized techniques as we have presented in this study to determine the extent to which ions are a controlling factor on their ecological range. Additionally, assessing the influence of more complex mixtures of inorganic ions and pH gradients on phage infection using high-throughput approaches as we have developed in this study will help characterize the complex higher-order interactions between geochemical context and phage:host dynamics.

### A conceptual model of spatial and temporal constraints on phage dynamics in terrestrial environments

Our suggestion that most freshwater environments are sub-optimal for successful phage infection (**Figure 3**) is ostensibly at odds with the prevalence of phage in some terrestrial soils [51] and lakes [8, 52]. However, in these environments, there are locations and times where ion concentrations locally or transiently will exceed the thresholds required for successful phage:host interactions. We propose that these “hot spots” and “hot moments” are critical sites of phage infection and should be considered in our conceptual models of phage ecology (**Figure 4, Figure S3**).

**Figure 4.**
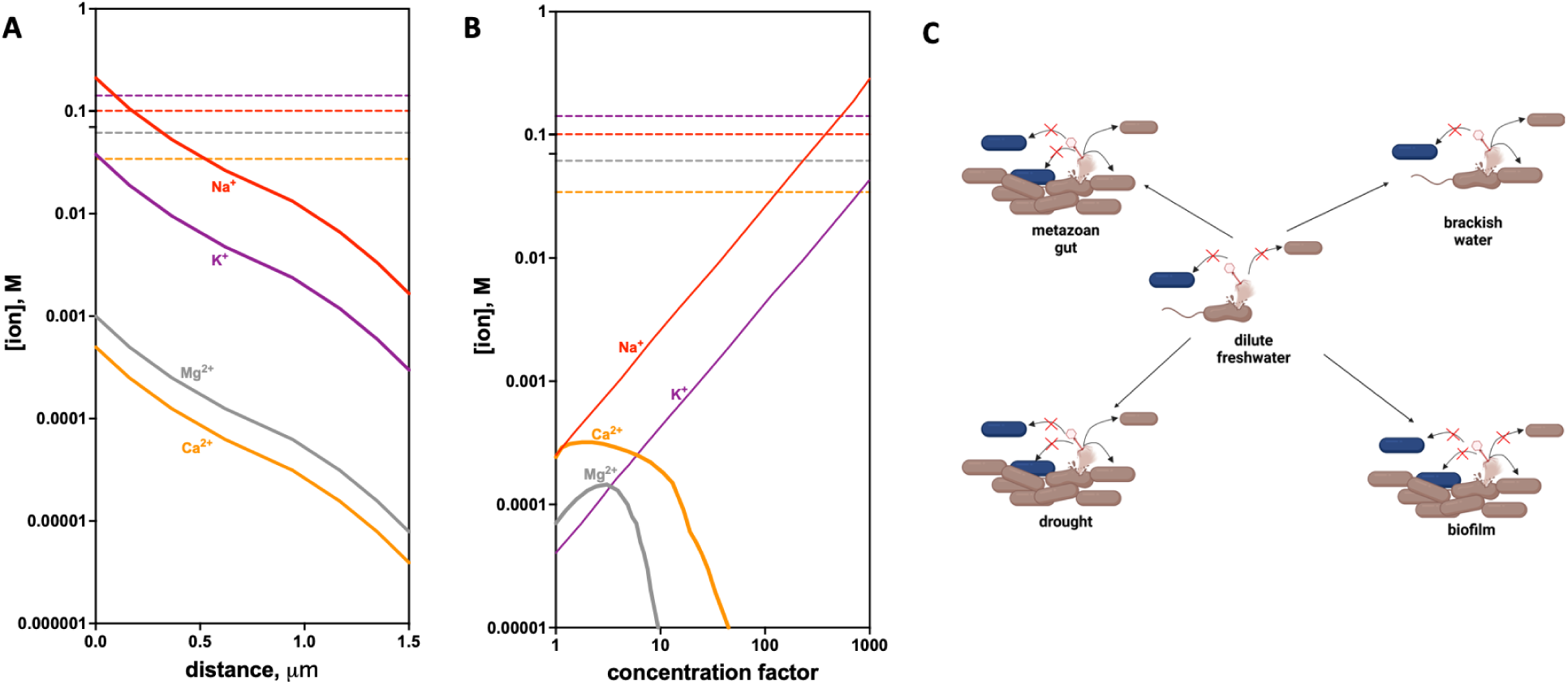
Temporal and spatial control of phage infectivity by ion thresholds in terrestrial ecosystems. **A.** Major cytoplasmic ion concentrations as a function of diffusion distance from lysed *E. coli* cells compared with EC_50_ concentrations required for T4 phage infection (dashed lines) **B.** Major ion concentrations in a typical Sierra Nevada spring water during evaporation as a function of concentration factor compared with phage EC_50_. Concentration factor is the ratio of the initial volume of water to the volume remaining after evaporation. Colors are consistent between EC_50_ and ion concentration lines. Colors and symbols are the same as in panel A. **C.** Conceptual model for phage infection. Ionic strength in freshwater environments is too low to promote phage infection. Higher ionic strength is found in metazoan guts or brackish/marine waters and transiently observed in drought influenced soils or in a biofilm adjacent to a lysed cell. When other strains (blue) are present, such as when the dominant electron donor/carbon source or other niche dimensions shift, fewer productive phage:host interactions will proceed.

In bacterial biofilms, diffusion of ions from the cytoplasm of bacterial cells lysed by phage will locally maintain sufficient ionic strength for infection of new hosts. We can model the diffusion of these ions from cells (**Figure 4A**). We know the ion concentrations in bacterial cytoplasms are ∼212 mM Na^+^, 38 mM K^+^, 0.5 mM Ca^2+^, 1 mM Mg^2+^ and a cell can be approximated as a sphere with a radius of one micron. Thus, we can calculate how these concentrations will decrease if they diffuse evenly into larger volumes. Our analysis shows that beyond ∼1 micron distance from lysed cells, the concentration of Na^+^ will drop below the EC_50_ for successful lytic infection by T4 phage (**Figure 4A**). This suggests that phage blooms in biofilms are propagated by very local diffusion dynamics between adjacent cells. Interestingly, T4 phage begins to aggregate at concentrations near the lytic infection EC_50_ for Na^+^ [17]. As phage aggregates are more stable to environmental stressors, this may suggest a mechanism phage have evolved to survive outside of bacterial hosts at low ionic strength when infection is disfavored.

During drought, ionic solutes in soil pore water are concentrated until the solubility constants (Ksp) for insoluble inorganic compounds are exceeded (**Figure 4B**). In many freshwater systems, the formation of magnesium silicates and calcium carbonates limit solubility of Ca^2+^ and Mg^2+^. Na^+^ and K^+^ salts are much more soluble and as such, the concentration of these ions will increase linearly with the concentration factor (Concentration factor is the ratio of the initial volume of water to the volume remaining after evaporation. A concentration factor of 3 means that ⅓ of the original water remains). Ion concentrations in the evaporation of a typical spring water (Sierra Nevada mountain springwater, CA) have been modeled [53] and when we overlay T4 phage EC_50_ on the ion concentration profiles we observe that between 100-1000 fold concentration of this water is necessary for phage to infect.

Thus, from our results and literature precedent, we propose a model in which phage infection proceeds when ionic strength is locally or transiently elevated (**Figure 4C**). Some recent field observations are consistent with our model. Metagenomic surveys of soils suggest a strong influence of soil moisture on viral ecological dynamics [54–56]. For example, a strong distance-decay relationship was observed in similarity between viral populations across a grassland field site [54]. This could be explained by the strong limitations on viral dispersal we anticipate when freshwater ion concentrations are too low to enable infection. Other studies have observed a strong influence of soil moisture content on the viral community content [55], but this may reflect the well established influence of drought on microbial community composition [57, 58]. Other field data support our model implicating a direct role of drought in controlling phage infection. Recently, in a timecourse from a soil undergoing rewetting after a period of drought, phage DNA was observed in the first hours after a rain event [59]. Based on our observations, this rapid phage bloom can best be explained by phage infection occurring during the early phases of the drought when host metabolism is slowing, with phage replication happening upon rewetting facilitated by the revival of host bacterial metabolism. Careful fieldwork to measure soil porewater ion concentrations and phage:host dynamics over a complete wet-dry-wet cycle would be helpful to further support this hypothesis.

Another feature of our conceptual model of phage predation is that the enrichment of non-host bacteria will decrease the probability of productive phage:host interactions. Our experiments with varying carbon sources provide support for this model. We observed that when *Sulfurospirillum* or *Klebsiella* were enriched in our cultures the predatory efficiency of the *Escherichia* and *Citrobacter* phage was decreased as well as the phage impact on DNRA activity. It follows that prebiotic nutrient amendments are potentially important to enhance the efficacy of phage cocktails, and that as carbon complexity increases (e.g. in soils) phage predation is likely to be less important in mediating microbiome composition and function. We anticipate that further work to assess how nutrient complexity influences phage:host interactions in natural systems will be important to support this hypothesis.

## Conclusions

Understanding the environmental controls on phage ecology is essential for a predictive understanding of the impact of the viral fraction on biogeochemical cycling in terrestrial ecosystems. Furthermore, as there is optimism about the use of phage and other genetic editing technologies to control and manipulate microbial ecosystems, it is important to consider the conditions under which phage-based microbiome engineering will be successful. With improvements in our ability to simulate the higher order interactions between environmental parameters we can add important constraints to our models of microbial dynamics in complex terrestrial systems. We anticipate that the use of low complexity model microbiomes that are archived, genomically characterized and manipulable in laboratories in high-throughput will yield important insights into how selective pressures influence microbiome structure and element cycling function.

## Materials and Methods

### Cultivation of bacteria in the presence of phage and inorganic ions and dose-response analysis

To quantify the influence of inorganic ions on the growth of *E. coli* in the presence of T4 phage we used a low ionic strength culture media containing 30 mM PIPES buffer pH 7, 5 mM ammonium chloride, 1 mM sodium phosphate, 1 mM L-cysteine, 30 mM D-glucose and DL vitamins and minerals. All chemicals are from Sigma-Aldrich (St Louis, Mo, USA). Final Na^+^ concentration in this media was ∼30 mM. In this culture media, phage were unable to infect and lyse *E. coli* in control cultures. All buffer and nutrient stock solutions were prepared with ultrapure water in plastic labware to minimize ion contamination from glassware. Cultures of *E. coli* recovered in LB were washed thrice in 2x concentrated medium prior to inoculation into growth assays. T4 and MS2 phage purified from *E. coli* LB cultures was dialyzed using Slidealyzer cassettes (Pierce, Thermo-Fisher, Waltham, MA, USA) into 30 mM PIPES buffer pH 7 to remove residual ions. Phage was mixed with *E. coli* cultures at a multiplicity of infection (MOI) of ∼1 (10^8^ phage per 10^8^ bacteria) in 2x concentrated growth medium. 40 μL of phage and *E. coli* in 2x media were then transferred into 384 well microplates (Costar, Thermo Fisher Scientific, Waltham, MA, USA) containing 40 μL of serially-diluted aqueous solutions of inorganic ion [30, 31]. Cultures were incubated at 30 °C and shaken at 700 rpm in a multitron shaker/incubator (Infors, Annapolis Jn, MD, USA). After 12 hours optical density was measured for dose-response analysis where normalized growth data was fit to a non-linear regression curve using GraphPad Prism (Graphpad software, Boston, MA, USA) to determine the half-maximal inhibitory concentrations for growth inhibition of the *E. coli* due to phage infection (EC_50_) and inorganic ion toxicity (IC_50_).

### Phage isolation and characterization

Phages used in this study are listed in Figure S1. We enriched lab stock of T4 and MS2 phages on BW25113 and C3000 *E. coli* strains respectively using standard protocols [60]. All other phages were isolated from local wastewater samples (East Bay Municipal Utility District, Berkeley, CA) using *E. coli* and *Citrobacter* isolates as hosts following the standard phage isolation protocol [61]. Phage titer was estimated by spotting 10-fold serial dilution of each phage in SM buffer (Teknova) on a lawn of target host using top agar overlay method with 0.7% LB agar. For phage dilutions in plaque assays we used an SM buffer supplemented with 10 mM calcium chloride and magnesium sulphate (Sigma-Aldrich). We routinely stored phages as filter-sterilised (0.22 μm) lysates at 4 °C. For genome sequencing, phage genome was extracted using Wizard genomic DNA purification kit (Promega, Madison, WI) and sequenced at Massachusets General Hospital next generation sequencing core facility. Genomes of both phage and bacteria used in this study are part of NCBI BioProject Accession PRJNA576510.

For transmission electron microscopy imaging, 3 µl of isolated bacteriophages were applied on a 300-mesh ultra-light carbon-coated copper grid for 5 min. The grid was briefly washed with sterile water and blotted with filter paper before staining with 3 µl of 2% aqueous uranyl-acetate. After 10 sec the grid was blotted to dryness. Preparations were examined with a 120KV Jeol1400-FLASH transmission electron microscope at Lawrence Berkeley National Laboratory. The sample was imaged at a magnification range of 12 and 50kX with the Oneview 16-Megapixel camera (Gatan®).

### Cultivation of freshwater nitrate reducing enrichment culture in the presence and absence of phage and inorganic ions

Freshwater nitrate reducing enrichment cultures were recovered from glycerol stocks in anoxic chemically defined basal medium supplemented with 2 grams/liter yeast extract as the sole organic carbon source and electron donor and 20 mM sodium nitrate as the sole terminal electron acceptor as previously described [14]. Basal medium contained per liter: 1.5 g sodium chloride, 2.5 g ammonium chloride, 10 g sodium phosphate, 1 g potassium chloride and 30 mM HEPES buffer with vitamins and minerals added from 100x stock solutions. Vitamin stock solution contained per liter: 10 mg pyridoxine HCl, 5 mg 4-aminobenzoic acid, 5 mg lipoic acid, 5 mg nicotinic acid, 5 mg riboflavin, 5 mg thiamine HCl, 5 mg calcium D,L-pantothenate, 2 mg biotin, 2 mg folic acid, 0.1 mg cyanocobalamin. Mineral stock solution contained per liter: 3 g magnesium sulfate heptahydrate, 1.5 g nitrilotriacetic acid, 1 g sodium chloride, 0.5291 g manganese(II) chloride tetrahydrate, 0.05458 g cobalt chloride, 0.1 g zinc sulfate heptahydrate, 0.1 g calcium chloride dihydrate, 0.07153 g iron(II) chloride tetrahydrate, 0.02765 g nickel(II) sulfate hexahydrate, 0.02 g aluminum potassium sulfate dodecahydrate, 0.00683 g copper(II) chloride dihydrate, 0.01 g boric acid, 0.01 g sodium molybdate dihydrate, 0.000197 g sodium selenite pentahydrate. All chemicals are from Sigma-Aldrich.

To measure the influence of carbon sources on the end-products of the archived nitrate reducing microbial communities, cells from recovered enrichment cultures were pelleted at 4000 RCF and washed three times with 2x concentrated basal medium lacking a carbon source. Washed cells were resuspended in 2x concentrated basal medium lacking a carbon source to an optical density (OD 600) of 0.04 and the cell suspension was transferred into either 384 well microplates (Costar) or 96 deep-well blocks (Costar) in which 94 carbon sources and water controls were arrayed (Table S1). Carbon source stock solutions were added to microplates using a Biomek FxP liquid handling robot (Beckman Coulter, Indianapolis, IN, USA) and kept in an anaerobic chamber (Coy, Coy Lab Products, Grass Lake, MI, USA) for 48 hours to become anoxic prior to inoculation using an Avidien Micropro 200 pipettor (Mettler-Toledo, Columbus, OH USA). Inoculated microplates were sealed with silicon microplate seals (VWR, Radnor, PA) and incubated at 30 °C in an incubator in an anaerobic chamber (COY). Growth was monitored by optical density (OD 600) using a Tecan M1000 Pro microplate reader (Tecan Group Ltd., Männendorf, Switzerland) and cultures were harvested at 48 hours for DNA sequencing and colorimetric assays to measure ammonium.

To measure the influence of inorganic ions on the efficacy of the phage cocktail, phage were dialyzed into low ionic strength buffer as described above and mixed with the washed enrichment culture and cultures containing 20 mM D-trehalose and 20 mM sodium nitrate in the presence of serially-diluted solutions of calcium chloride or sodium chloride. As above, culture were incubated for 48 hrs prior to measurements of optical density and ammonium

### Colorimetric assays to measure ammonium production in nitrate-reducing enrichment cultures

Ammonium production was measured as previously described [14]. Microplate seals were removed from 384-well microplates containing enrichment cultures and a Biomek FxP (Beckman Coulter) was used to transfer 4 µL of culture to assay microplates prefilled with 20 µL of distilled deionized. In sequential order, 4 µL of citrate reagent, 8 µL of salicylate/nitroprusside reagent, and 4 µL bleach reagent were added to assay plates which were then kept at 30 °C for 30 min. Citrate reagent contains 10 g trisodium citrate and 4 g sodium hydroxide in 200 mL water. Salicylate/nitroprusside reagent contains 15.626 g sodium salicylate and 0.250 g sodium nitroprusside in 200 mL water at pH 6–7. Bleach reagent contains 1 g sodium phosphate monobasic, 2 mL 2 M sodium hydroxide, 10 mL bleach (0.7 M NaOCl, Chlorox Company, Pleasanton, CA, USA) in 100 mL water at pH 12–13. For all colorimetric assays, we confirmed that interference of media additives (carbon sources and inorganic ions) was negligible. Using constants obtained from the BioNumbers database [62], we estimated the quantity of nitrogen assimilated into biomass by assuming 0.3 g/L of dry weight of bacterial culture at OD 600 = 1 [63, 64], and by assuming 12% nitrogen by weight in microbial biomass based on measured C:N:P ratios [65, 66].

### 16S rDNA amplicon sequencing of microbial enrichments to identify microbial community shifts associated with changes in element cycling end-products

For DNA extraction microbial cells from 500 µL cultures were pelleted by centrifugation at 4000 RCF after 48 hours of growth at 30°C. DNA (gDNA) extractions were performed as follows. Cell pellets were resuspended in 180 µL enzymatic lysis buffer (ELB) which contains 20 mM Tris-HCl pH 8.0, 1 mM sodium EDTA (TE) with 1.2% Triton X-100. 20 #x1D707;L lyzosyme (25 mg/mL) was added and the samples incubated overnight at 37 °C. The following morning 20 #x1D707;L of proteinase K (20 mg/mL) and 200 µL 4M guanidinium HCL were added and the samples incubated overnight at 55 °C. The following morning 4 µL RNAse A (100 mg/mL) was added and samples incubated for 2 hrs at room temperature. 200 µL of ethanol was added to precipitate DNA and samples were then applied to 96 well silica membrane plates (BPI-Tech, San Diego, CA, USA). DNA bound to membranes was washed twice with 400 µL 70% ethanol:TE and eluted in 100 µL TE. An Avidien Micropro 200 (Avidien) was used to transfer large volumes of buffers and multichannel pipettes (Rainin) used to apply samples to 96 well columns and add low volume reagents. A vacuum manifold (Qiagen, Redwood City, CA, USA) was used to perform column purification steps. Extractions using this method were found to be similar in terms of DNA yield and recovery to using the Gene QIAamp 96 DNA QIAcube HT Kit (Qiagen).

gDNA template was added to a PCR reaction to amplify the V4/V5 16S gene region using the 515F/ 926R primers based on the Earth Microbiome Project primers [67, 68] but with in-line dual Illumina indexes [69, 70]. The amplicons were sequenced on an Illumina MiSeq (Illumina, San Diego, CA, USA) at the QB3 Genomics facility (QB3 Genomics, UC Berkeley, Berkeley, CA, RRID:SCR_022170) with 2×300 bp Ilumina v3 reagents. Reads were processed with custom Perl scripts implementing Pear for read merging [71], USearch [72] for filtering reads with more than one expected error, demultiplexing using inline indexes and Unoise [73] for filtering rare reads and chimeras. 16S sequences in the relative abundance table were searched against the RDP database [74] to assign taxonomy. The functional capacity for nitrate reduction enzymatic steps was determined in a previous publication [14]. The closest 16S rDNA ESVs from 16S taxonomy were matched to the GTDB-Tk [75] taxonomic bins [14].

## Data Availability

DNA sequencing data are available under BioProject Accession PRJNA576510.

## Supplementary Tables and Figures

**Table S1.** Inhibitory potencies of 80 inorganic ion array in the absence of phage (IC50) or presence of phage (EC50). All measurements are in molar units (M) and 95% confidence intervals are reported.

**Figure S1.**
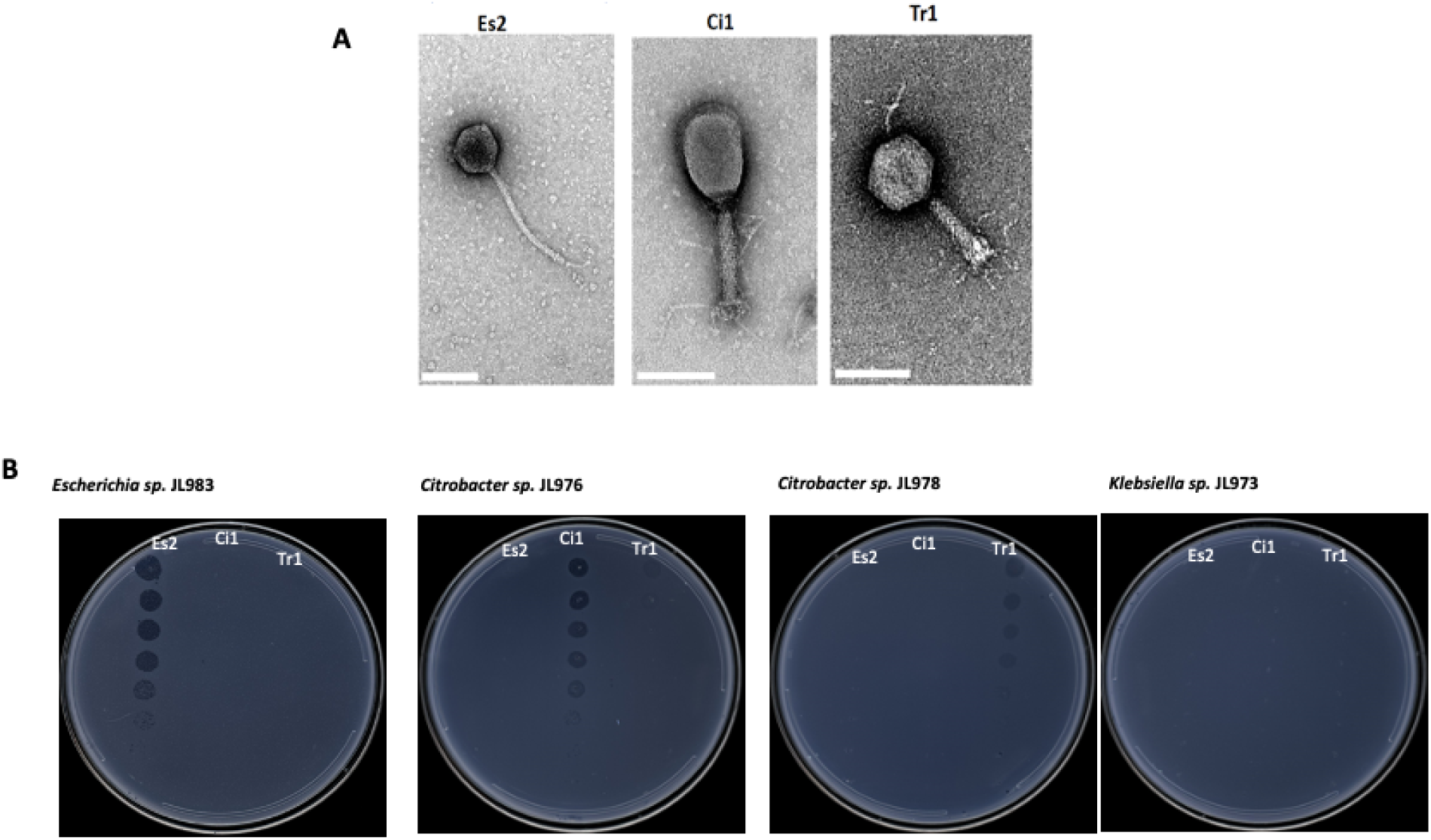
Phages in cocktail formulated to inhibit DNRA in freshwater nitrate reducing enrichment (FN) cultures. **A.** Transmission electron micrographs of ES2, Ci1 and Tr1 phages. The scale bar denotes 100 nm **B.** Plaque assays demonstrating specificity of phages for dominant *Enterobacteria* in the FN culture. ES2 is specific for *Escherichia* JL 983, Ci1 is specific for *Citrobacter sp.* JL 976 and Tr1 is specific for *Citrobacter sp.* JL 978.

**Figure S2.**
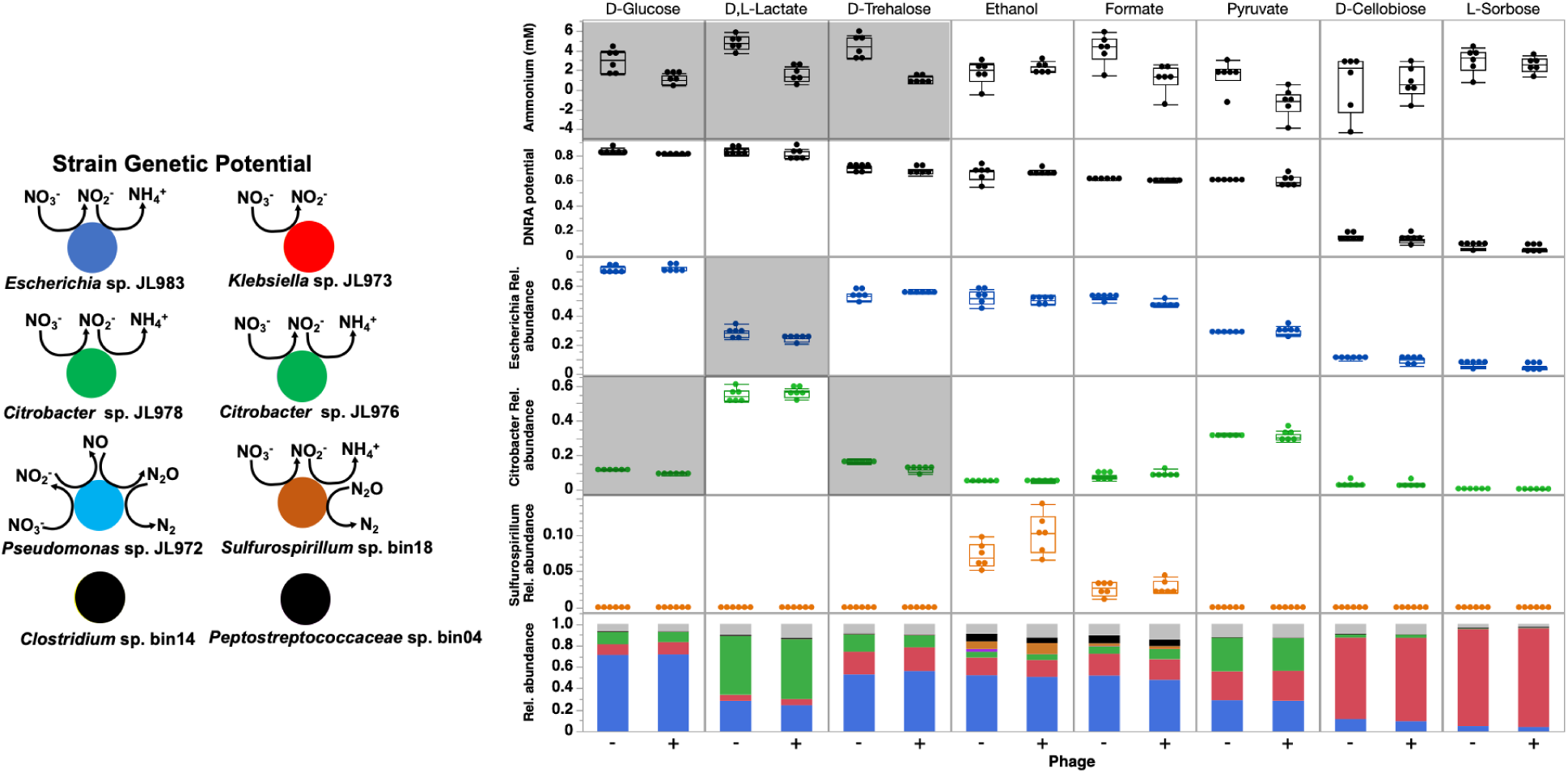
Ammonium production, DNRA genetic potential and relative abundances of dominant strains in the FN microbiome on selected carbon sources. In box plots: Box and whiskers represent interquartile range. Shaded panels indicate significant differences (ANOVA, p<0.05) between minus (-) and plus (+) phage conditions. In stacked bar plot: Dark Blue = *Escherichia*, Green = *Citrobacter*, Red = *Klebsiella*, Orange = *Sulfurospirillum*, light blue = *Pseudomonas,* Black = Gram-positive fermenters, Gray = other strains.

**Figure S3.**
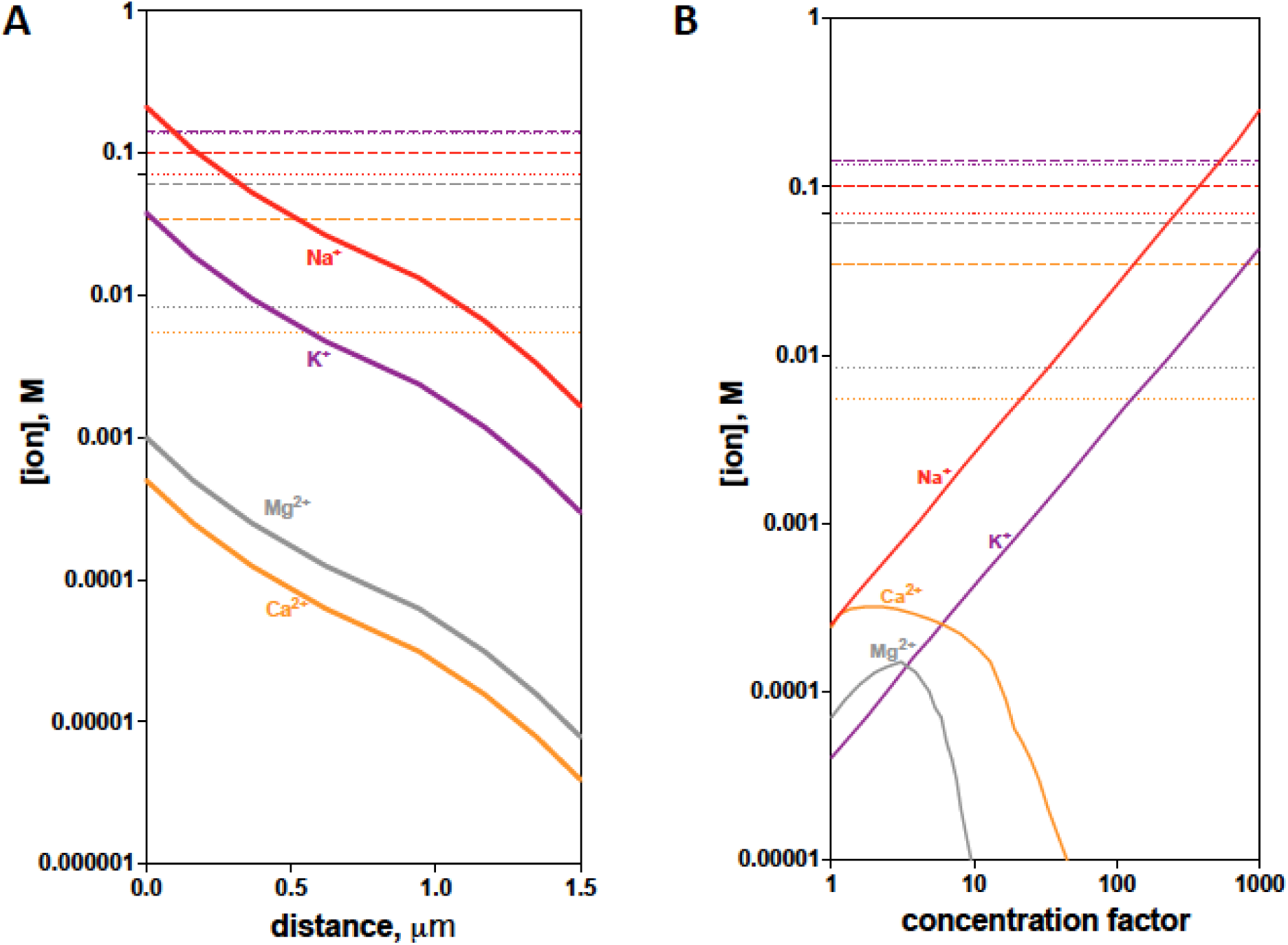
Temporal and spatial control of phage infectivity by ion thresholds in terrestrial ecosystems. **A.** Major cytoplasmic ion concentrations as a function of diffusion distance from lysed *E. coli* cells compared with EC_50_ concentrations required for T4 (dash), or MS2 (dotted) phage infection. **B.** Major ion concentrations in a typical Sierra Nevada spring water during evaporation as a function of concentration factor compared with phage EC_50_. Colors are consistent between EC_50_ and ion concentration lines. (colors and symbols are the same as panel A).

